# Single-cell multiomics reveals disrupted glial gene regulatory programs in Alzheimer’s disease via interpretable machine learning

**DOI:** 10.1101/2025.03.14.643349

**Authors:** Shuwen Zhang, Hongru Hu, Xue Wang, Chengjie Xiong, Yan W Asmann, Yingxue Ren

## Abstract

Recent development of single-cell technology across multiple omics platforms has provided new ways to obtain holistic views of cells to study disease pathobiology. Alzheimer’s disease (AD) is the most common form of dementia worldwide, yet the detailed understanding of its cellular and molecular mechanisms remains limited. In this study, we analyzed paired single-cell transcriptomic (scRNA-seq) and chromatin accessibility (scATAC-seq) data from the Seattle Alzheimer’s Disease Brain Cell Atlas (SEA-AD) Consortium to investigate the molecular mechanisms of AD at a cell-subpopulation-specific resolution focusing on glial cells. We benchmarked various multi-omics integration methods using diverse metrics and built an analytic workflow that enabled effective batch correction and cross-modality alignment, creating a unified cell state space. Through integrative analysis of 26 human brain samples, we uncovered AD-associated gene expression and pathway changes in glial subpopulations and highlighted important transcriptomic and epigenomic signatures via functional inference and interpretable machine learning paradigms, discovering the profound involvement of the Solute Carrier proteins (SLC) family genes in multiple glial cell types. We also identified glial cell-specific regulatory programs mediated by key transcription factors such as *JUN* and *FOSL2* in astrocytes, the Zinc Finger (ZNF) family genes in microglia, and the SOX family of transcription factors in oligodendrocytes. Our study provides a comprehensive workflow and a high-resolution view of how glial regulatory programs are disrupted in AD. Our findings offer novel insights into disease-related changes in gene regulation and suggest potential targets for further research and therapy.

## INTRODUCTION

Among the diverse cell populations in the brain, glial cells play essential roles in maintaining homeostasis, supporting neuronal function, and responding to injury (1) (2). Astrocytes, the most abundant glial cells, regulate synaptic activity, and provide metabolic support (3). Microglia serve as the resident immune cells of the brain, continuously surveying the environment and mounting immune responses to pathogens and cellular debris (4). Oligodendrocytes produce myelin, which insulates axons and facilitates efficient signal transmission (5), while oligodendrocyte precursor cells (OPCs) contribute to myelin repair and neural plasticity (6).

Growing evidence suggests that glial dysfunction is a key driver of neurodegenerative diseases, including Alzheimer’s disease (AD) (7,8), yet the precise regulatory mechanisms underlying these changes remain poorly understood. Investigating glial-specific gene regulatory programs could provide critical insights into how these cells contribute to AD pathology.

Understanding complex neurodegenerative diseases at the cellular level requires dissecting both gene expression programs and their regulatory mechanisms. While single-cell RNA sequencing (scRNA-seq) reveals the transcriptional landscape of individual cells (9), single-cell Assay for Transposase-Accessible Chromatin (scATAC-seq) uncovers chromatin accessibility patterns that shape gene activity (10,11). Together, these technologies, especially the recent joint profiling assays (12) (13), provide holistic views of cell states and gene regulation, where researchers are able to trace how transcription factors, enhancers, and promoters coordinate cellular identities and functions (11,14). However, integrating these datasets is challenging due to differences in data structure, feature types, sparsity, effects from technical batches, and the difficulty of directly linking chromatin accessibility to gene expression across individual cells. Unlike scRNA-seq, which has been long adopted by researchers with clear best practices for downstream analysis (15,16), multiomic data lacks standardized workflows, making it difficult to extract biologically meaningful insights.

To address these challenges, we leveraged the paired single-cell multiomic (scMultiome) data from the Seattle Alzheimer’s Disease Brain Cell Atlas (SEA-AD) Consortium and developed a comprehensive workflow for analysis (**Figure 1**). SEA-AD is a large-scale initiative that systematically profiles gene expression and chromatin accessibility across different brain regions in AD and healthy individuals (17). Our analysis focused on the middle temporal gyrus (MTG), a cortical region that is among the earliest affected in AD and is critical for language processing and episodic memory (18) (**Figure 1a**). By applying our workflow on SEA-AD datasets, we aimed to uncover the molecular alterations driving AD-related changes in glial cells within this brain region.

**Figure 1.**
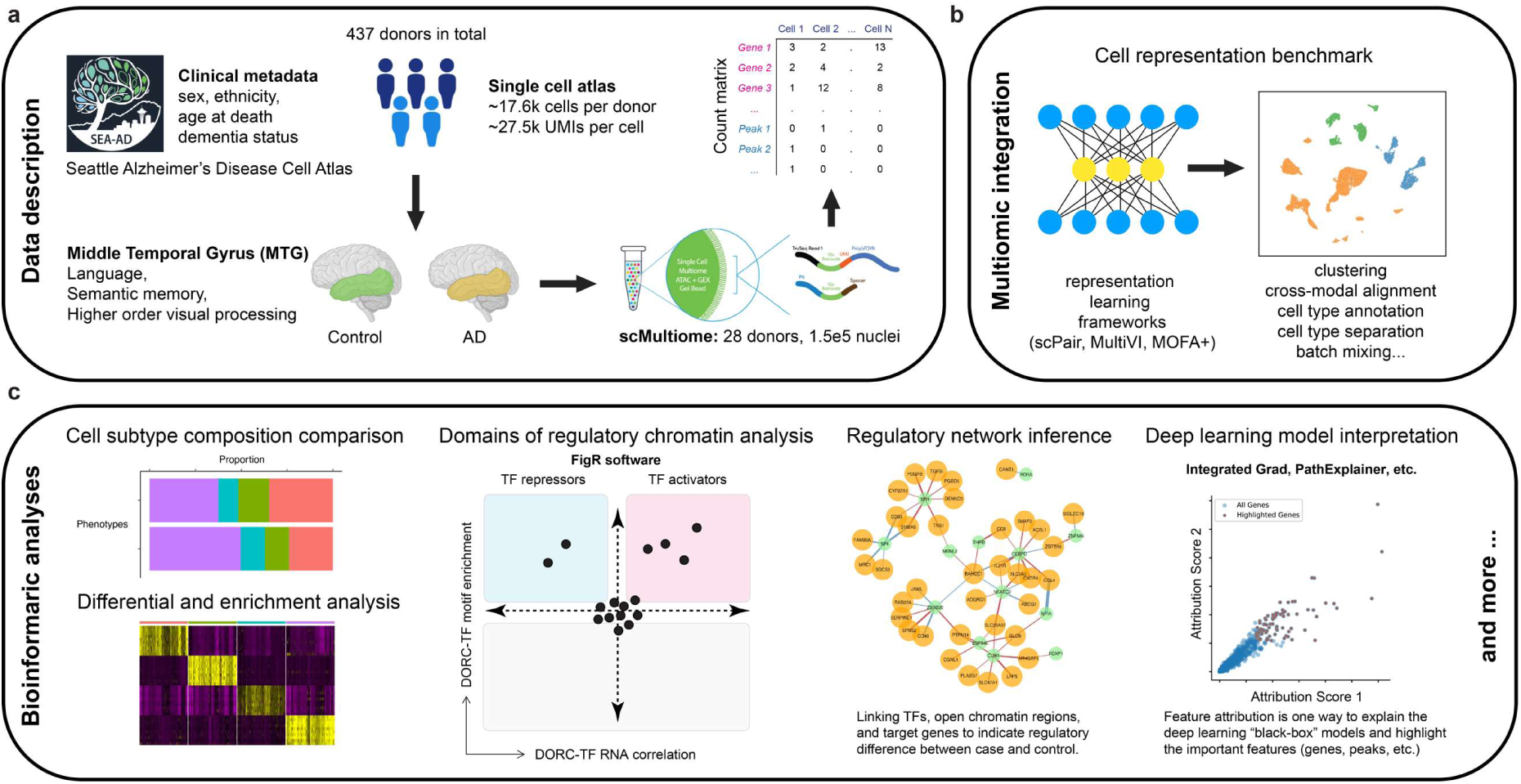
Overview of SEA-AD multiome dataset, integration tool benchmarking, and downstream analyses. (**a**) The SEA-AD dataset comprises comprehensive multimodal data, and this study specifically focused on scMultiome data from the middle temporal gyrus (MTG) that profiles both transcriptomic (RNA) and chromatin accessibility (ATAC) modalities across various cellular states. (**b**) A schematic of the multimodal cell representation learning benchmarking, emphasizing the evaluation tasks and integration performance assessment across diverse tools designed to handle paired single cell multiomics data. (**c**) Key downstream analyses include cell subtype composition and proportion comparison, differential and enrichment analysis, Domains of Regulatory Chromatin (DORC) Analysis, regulatory network inference, and model interpretation using feature attribution technologies.

Within the workflow, we considered an effective integration strategy as a key initial step to better capture cellular heterogeneity and regulatory dynamics in AD. Multiomic integration not only helps account for technical variations such as batch effects and donor-specific differences, but also provides a more holistic view of cell states by aligning gene expression with chromatin accessibility. Proper alignment of these modalities ensures that meaningful biological signals are preserved, enabling a more accurate characterization of cell types and their regulatory landscapes. However, due to differences in data structure and modality-specific noise, selecting the most effective integration method remains a critical challenge. To address this, we systematically benchmarked multiple computational approaches for integration by comparing factorization-based models and deep learning methods to determine the most effective strategy for aligning scRNA-seq and scATAC-seq data (**Figure 1b**). Factorization-based models, such as MOFA+ (Multi-Omics Factor Analysis) (19,20), decompose multiomic data into shared factors, making them useful for identifying biologically relevant variation. Deep learning-based approaches introduce more flexible, data-driven strategies for aligning different modalities.

Among these, MultiVI (21) applies unsupervised variational autoencoders for cross-modality alignment, while scPair (22) implements a supervised learning approach that links chromatin accessibility and gene expression end to end. These frameworks are powerful for learning biologically meaningful low-dimensional representations of the data across modalities.

For the downstream analyses from our workflow, we first leveraged the learned cell state space from the most suitable method to identify differential gene expression and chromatin accessibility patterns, and highlighted key molecular signatures that drove disease-associated changes. Afterwards, we reconstructed gene regulatory programs for different glial subpopulations to demonstrate how transcription factor activity shifts in AD. Methods such as figR (23), which has been applied for functional inference based on cis-regulatory associations (gene-peak correlations), helped establish putative gene regulatory networks and identify critical marker genes and domains of regulatory chromatin (DORCs). Beyond these standard analyses steps, we applied interpretable machine learning techniques to analyze the embeddings learned by the selected cell state inference model in order to validate these computational findings *in silico* (**Figure 1c**). This approach allowed us to identify genes that strongly contribute to integrated cell states, helping pinpoint key regulators of glial dysfunction in AD.

Taken together, our study provides an optimized framework for single-cell multiomic integration and analysis while demonstrating the value of interpretable machine learning in cross-validating computational findings and uncovering key molecular signatures. This analysis offers a clearer view of the molecular signatures driving glial dysfunction in AD, providing insights into the regulatory mechanisms underlying glial contributions to disease pathology.

## MATERIAL AND METHODS

### Dataset description

We obtained the Seattle Alzheimer’s Disease Brain Cell Atlas (SEA-AD) dataset for the middle temporal gyrus (MTG) from two sources (accessed September 25, 2023): the multiome data files from Synapse (ID: syn29879329) and the processed RNA expression matrix from Chan Zuckerberg CELL x GENE (https://cellxgene.cziscience.com/collections/1ca90a2d-2943-483d-b678-b809bf464c30). The initial dataset contained 98,601 cells from 16 normal donors and 52,755 cells from 12 dementia donors.

### Sample selection and data processing for scMultiome integration benchmarking

To benchmark scMultiome integration methods, we selected four donors to ensure balanced representation across conditions and biological sexes while maximizing cell numbers. These donors included H20.33.016 (female with dementia, 8,130 cells), H20.33.017 (male with dementia, 6,797 cells), H20.33.008 (female normal, 8,414 cells), and H20.33.025 (male normal, 8,950 cells).

For scATAC-seq data processing, we first conducted peak calling using MACS2 on ATAC fragments from these four donors using the HG38 reference genome. After excluding non-autosome peaks, we identified 190,133 peaks. We then filtered peak regions opening in less than 1% of cells per individual and kept the intersection set across all four donors, resulting in 151,859 peak features across 32,291 cells. We further filtered cells to include only those with 300-30,000 peak features, resulting in 30,090 cells for downstream analysis. To process scRNA-seq data, we removed genes present in less than 1% of cells per individual. Taking the intersection set across all four donors yielded 19,400 gene features. We then filtered cells to include only those with 1,000-150,000 gene features, resulting in 32,289 cells for downstream analysis.

### Benchmarking sc-multiome integration methods

We collected the intersection of 30,089 cells present in both data modalities, and split them into a modeling set containing 24,072 cells (80%) and a held-out test set of 6,017 cells (20%). The modeling set was further divided into 19,258 cells (80%) for training and 4,814 (20%) cells for validation to enable early stopping and prevent overfitting. To meet modeling requirements using MultiVI and scPair, we converted ATAC peak counts to binary values.

For scPair execution, we followed the tutorial guidelines available on GitHub (https://github.com/quon-titative-biology/scPair/tree/main/tutorials). The ATAC to RNA component utilized two hidden layers with 900 and 30 nodes, respectively, and a latent dimension of 30. We set the learning rate to 0.001, batch size to 300, and employed batch normalization and layer normalization with GELU activation function. We used the Adam optimizer and set the dropout rate to 0.1, and. For the output, we implemented the zero-inflated negative binomial distribution (ZINB) for RNA counts. The RNA to ATAC component maintained similar settings but employed the Bernoulli distribution for the binarized peaks. Sex, donor ID, and disease status were included as covariates using one-hot encoding during model training. The mapping networks were implemented as linear transformations.

We ran MultiVI using scvi-tools and followed the original paper (24) by not explicitly incorporating the modality penalty term in the loss function. We designated donor ID as the “batch_key” and set the latent dimension as 30 while keeping other parameters at their default values as specified in the tutorial: https://docs.scvi-tools.org/en/stable/tutorials/notebooks/multimodal/MultiVI_tutorial.html.

For MOFA+ testing, we first preprocessed the data using the muon package (https://muon.scverse.org). The RNA modality underwent library size normalization with a size factor of 10,000, followed by log1p transformation for variance stabilization. We then selected highly variable genes using the setting of flavor=”seurat”, resulting in 3,606 genes. For the ATAC modality, we applied the “Term Frequency Inverse Document Frequency of records” (TF-IDF) transformation with a scale factor of 10,000 and performed feature selection using the setting of min_mean=0.05, max_mean=1.5, min_disp=0.5 as suggested in the tutorial: https://muon-tutorials.readthedocs.io/en/latest/single-cell-rna-atac/pbmc10k/2-Chromatin-Accessibility-Processing.html. We also utilized mofapy2 (https://github.com/bioFAM/mofapy2), with 30 factors (K=30), enabling ard_weights and ard_factors, and employed GPU acceleration with “fast” convergence mode to optimize processing time. Afterward, we utilized mofaX (https://github.com/bioFAM/mofax) to extract the variables of loadings and factors and fetch the values of 30-dimensional representations.

### Benchmark performance evaluation

To evaluate performances of the three integration methods, we first investigated cell representation quality using classical measures, including cell clustering annotation accuracy by Adjusted Rand Index (ARI) (25) and Normalized Mutual Information (NMI) (26), cell type separation by Average Silhouette Width (ASW)(27) and Cell-specific Local Inverse Simpson’s Index (cLISI) (28), and batch mixing by batch-specific Average Silhouette Width (batchASW) (29).

We further assessed cross-modality alignment using Fraction of Samples Closer than True Match (FOSCTTM)(30). For cross-modality prediction, we employed multiple metrics including Pearson Correlation Coefficient (PCC) and Spearman Correlation Coefficient (SCC) for RNA prediction, and Area Under ROC Curve (auROC) and 1-Binary Cross Entropy (1-BCE) for ATAC prediction.

### scMultiome integration for glial cells using scPair

We applied scPair on 26 of the 28 donors from the SEA-AD MTG multiome dataset focusing on glial cells, excluding H20.33.040 and H20.33.043 due to their relatively low glial cell counts (48 and 416 cells, respectively). The remaining donors each contributed more than 1,000 cells.

From scPair-learning embeddings, we obtained of 25,054 cells, 104,140 peaks, and 18,088 genes. We also performed separate peak calling for each glial cell types (microglia, astrocytes, oligodendrocytes and OPC) to ensure that we capture cell-type-specific peaks for cell-population-specific scPair modeling. The training settings were consistent with the benchmarking experiments.

### Data normalization, cell clustering, and differential expression analysis

Seurat (v5.0.0) (16) was employed for all downstream analyses, starting with global-scaling normalization using the ‘LogNormalize’ method, followed by log-transformation for downstream analyses. Highly variable features (default: 3,000 genes) were identified for each sample using the ‘FindVariableFeatures’ function to highlight biological signals. Linear transformation was applied to the combined dataset as a standard pre-processing step. Harmony (28) was used for batch correction and primary clustering based on scPair-derived representations. Principal component analysis (PCA) was conducted for dimensionality reduction, and the Uniform Manifold Approximation and Projection (UMAP) (31) method was used to visualize the integrated dataset. The ‘FindNeighbors’ and ‘FindClusters’ functions were used for cell clustering, with cluster-specific markers identified using the ‘FindConservedMarkers’ function. Clusters were annotated to known cell types based on marker genes. Differentially expressed genes (DEGs) between AD and controls were defined using a significance threshold of adjusted P < 0.05 and |log2 fold change| > 1. Visualization of marker genes and DEGs was conducted using the ‘FeaturePlot’ and ‘VlnPlot’ functions in Seurat alongside the ggplot2 package (https://ggplot2.tidyverse.org). The enrichR package (32) was used for functional enrichment analysis of DEGs.

### Glial cell subpopulation analysis and feature attribution calculations

Each glial cell types were re-clustered to reveal subpopulations using the same Seurat procedures as previously described. For scPair model interpretation, we employed two widely used feature attribution approaches in single cell studies: Integrated Gradients implemented by Captum package (https://captum.ai) (33) and Path-Explainer from PAUSE (https://github.com/suinleelab/PAUSE) (34). We used cluster-specific averaged input values as the baseline for feature attribution, calculating and averaging attribution scores within clusters. After selecting the top 100 highly attributed features per method per cluster, we identified highly-contributing features through intersection across methods and clusters.

### Chromatin opening visualization

For ATAC chromatin opening visualization on the genome, we used the function CoveragePlot() from Signac (https://stuartlab.org/signac/) (35) with selected chromatin regions and corresponding genes.

### Domain of Regulatory Chromatin (DORC) analysis

We employed the Functional Inference of Gene Regulation (FigR) (23) framework to interrogate the scMultiome data for the inference of gene regulatory networks (GRNs). Cis-regulatory associations were assessed by calculating gene-peak correlations, followed by the definition of Domains of Regulatory Chromatin (DORCs), representing loci where chromatin accessibility and gene expression are tightly coupled. The core regulatory interactions were then inferred through the FigR framework’s GRN inference function, which integrates DORCs with scRNA-seq data to identify transcription factors (TFs) likely to regulate target genes. The identified TF-gene pairs were filtered by statistical significance, and the resulting GRN was subjected to network analysis and functional enrichment.

## RESULTS

### A customized workflow for single cell multiomic analysis

To analyze single-cell data from the SEA-AD consortium, we developed a customized workflow for integrating and analyzing this multiomic data. The selected dataset includes over 150,000 nuclei from 28 donors, covering both Alzheimer’s disease (AD) and control samples and both biological sexes. We began by evaluating multimodal cell representation learning tools to determine the most suitable integration method. Our benchmarking included three representative integration frameworks: scPair (22), a supervised deep learning method; MultiVI (21), an unsupervised deep generative method; and MOFA+ (20), a factorization-based method. Based on the evaluation results, we selected the top-performing method and integrated it into our workflow for downstream bioinformatics analyses. These included cell subtype composition analysis, differential gene expression and enrichment analysis, Domains of Regulatory Chromatin (DORC) analysis, regulatory network inference, and feature attribution for model interpretation (**Figure 1c**).

### Benchmarking of scMultiome integration tools suggested top performer for proper integrative analysis

In recent years, several computational methods have been developed for integrating multiomic data, but their effectiveness varies depending on the dataset characteristics and analytical goals. To determine the most suitable tool, we compared scPair, MultiVI, and MOFA+, which represent three distinct integration approaches. Our evaluation was conducted on a subset dataset of four samples, selected based on high cell counts and balanced representation across biological sex and disease status. We first assessed cell state representation learning, focusing on key integration performance metrics such as clustering accuracy, cell separation, and batch effect correction (**Figure 2a**). Clustering accuracy was measured using Adjusted Rand Index (ARI) and Normalized Mutual Information (NMI), while cell type separation was evaluated using Average Silhouette Width (ASW) and cell-type Local Inverse Simpson’s Index (cLISI). To quantify batch effect removal or batch mixing while preserving biologically meaningful signals, we used batch-specific ASW and integration Local Inverse Simpson’s Index (iLISI).

**Figure 2.**
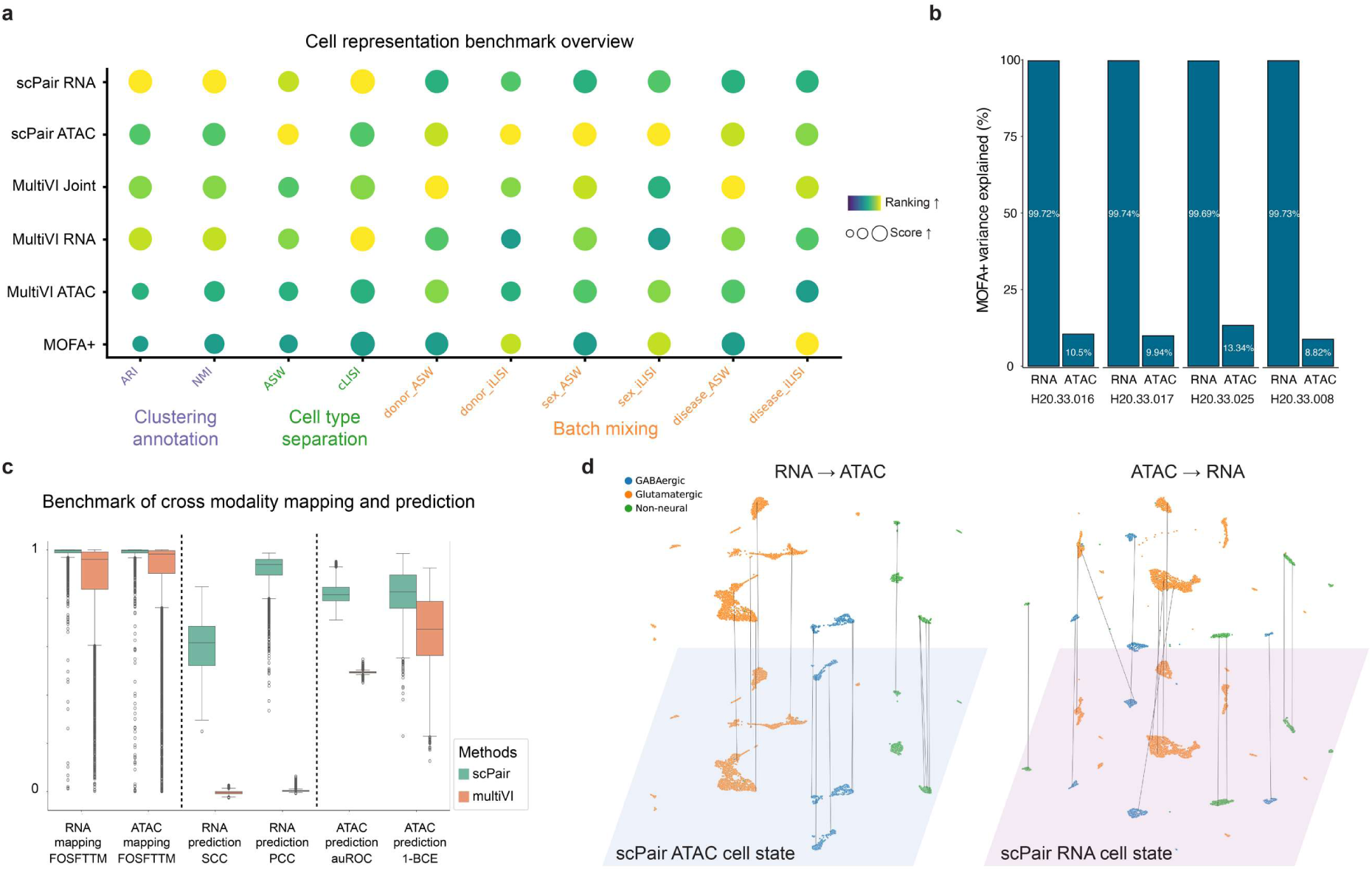
Benchmarking of scMultiome test data using scPair, MultiVI and MOFA+. (**a**) Evaluation of cell representation generated by scPair, MultiVI and MOFA+, displaying each method’s ranking and score across the different evaluation criteria. Methods are evaluated based on their capacity for clustering annotation (measured by ARI and NMI), cell type separation (ASW and cLISI), and batch mixing (covariate-specific ASW and iLISI metrics). Higher ranks and larger circle sizes indicate better performance. (**b**) Percentage of variance exampled by RNA and ATAC using MOFA+ in each of the four testing samples. (**c**) Performance benchmarking of scPair and MultiVI across cross-modality mapping tasks, with evaluation metrics calculated on held-out test sets. Box plots display performance metrics including 1 - Fraction of Samples Closer Than the True Match (1-FOSCTTM), Spearman Correlation Coefficient (SCC), Pearson Correlation Coefficient (PCC), area under the Receiver Operating Characteristic Curve (auROC), and 1 - Binary Cross Entropy (1-BCE), where higher values indicate better performance. In the box plots, the minima, maxima, centerline, bounds of box, and whiskers represent the minimum value in the data, maximum, median, upper and lower quartiles, and 1.5x interquartile range, respectively. (**d**) Left: UMAP visualization of the ATAC-based (ground truth, bottom) and RNA-to-ATAC (mapped, top) cell state spaces as learned by scPair using the held-out test set. Each point represents a cell, with lines connecting each cell’s observed ATAC state and mapped ATAC state (derived from RNA state). Colors indicate major cell classes based on original study annotations. Right: An analogous UMAP for RNA-based (ground truth, bottom) and ATAC-to-RNA (mapped, top) cell states, also learned by scPair using the same held-out test set.

Among the three methods, scPair ranked highest across the most of integration metrics, followed by MultiVI, with MOFA+ ranking lowest in most cases (**Figure 2a**). Although MOFA+ is designed to be an interpretable factorization method, its integration was biased toward RNA and led to underutilization of chromatin accessibility information (**Figure 2b**). In contrast, scPair and MultiVI generated separate embeddings for RNA and ATAC and have objectives of predicting or reconstructing the modality-specific features for each cell, which allows for a more balanced flow of modality-specific information.

Cross-modal alignment and prediction are also critical functionalities of a tool for linking gene expression (RNA) to chromatin accessibility (ATAC). Unlike scPair and MultiVI, MOFA+ lacks key capabilities such as modality-specific embedding inference and cross-modality imputation. This means that scPair and MultiVI are able to predict missing modality data after training while MOFA+ cannot. Given these limitations, we directly compared scPair and MultiVI in cross-modality alignment and prediction tasks, To evaluate alignment performance, we measured 1 - Fraction of Samples Closer Than the True Match (1-FOSCTTM) and found that scPair outperformed MultiVI (**Figure 2c**). Similarly, when assessing cross-modality prediction, scPair achieved higher Spearman Correlation Coefficient (SCC) and Pearson Correlation Coefficient (PCC) for ATAC-to-RNA predictions, and higher area under the Receiver Operating Characteristic Curve (auROC) and 1 - Binary Cross Entropy (1-BCE) for RNA-to-ATAC predictions (**Figure 2c**).

Overall, our results demonstrated that scPair was the most effective integration tool for this dataset, outperforming the alternatives in cell clustering, batch effect correction, and cross-modality prediction accuracy. Using scPair-learned embeddings, we successfully mapped cell states between RNA and ATAC, allowing us to clearly distinguish major neuronal subtypes (**Figure 2d**). This well-defined cell state mapping lays a strong foundation for further analyses of the complete SEA-AD scMultiome dataset. Based on these results, we selected scPair as the top-performing integration tool and applied it to the full scMultiome dataset and focusing on glial cell populations. After training the model, we generated integrated embeddings and proceeded with downstream analyses to investigate cell-type-specific disease mechanisms.

### Glial cell diversity and cell type-specific transcriptome dysregulation in Alzheimer’s middle temporal gyrus (MTG)

Historically, neurons were considered the primary signaling cells of the nervous system while glial cells were thought to serve the merely supportive role in maintaining neural architecture (36). However, accumulating evidence over the past decade has proven the critical functions of glial cells in both the structural and dynamic regulation of neural networks and Alzheimer’s disease pathobiology (8,37,38). As a proof of principle, we decided to focus on glial cells for both our workflow development and scientific investigation.

Using the best-performed integration method scPair, we analyzed the scMultiome data from 26 SEA-AD donors with > 1000 glial cells (**MATERIAL AND METHODS**). This approach successfully recaptured major glial cell types in the middle temporal gyrus (MTG) region, including astrocytes, microglia, oligodendrocytes and oligodendrocyte precursor cells (OPCs) (**Figure 3a**). These cell types were confirmed by the distinct expression patterns of classical marker genes, including *AQPR* for astrocytes, *CSF1R* for microglia, *MOBP* for oligodendrocytes, and *PDGFRA* for OPCs (**Figure 3b**). The chromatin accessibility profiles from the ATAC component further confirmed open chromatin regions near these marker genes (**Figure 3c**), which reinforces cell-type-specific regulatory landscapes.

**Figure 3.**
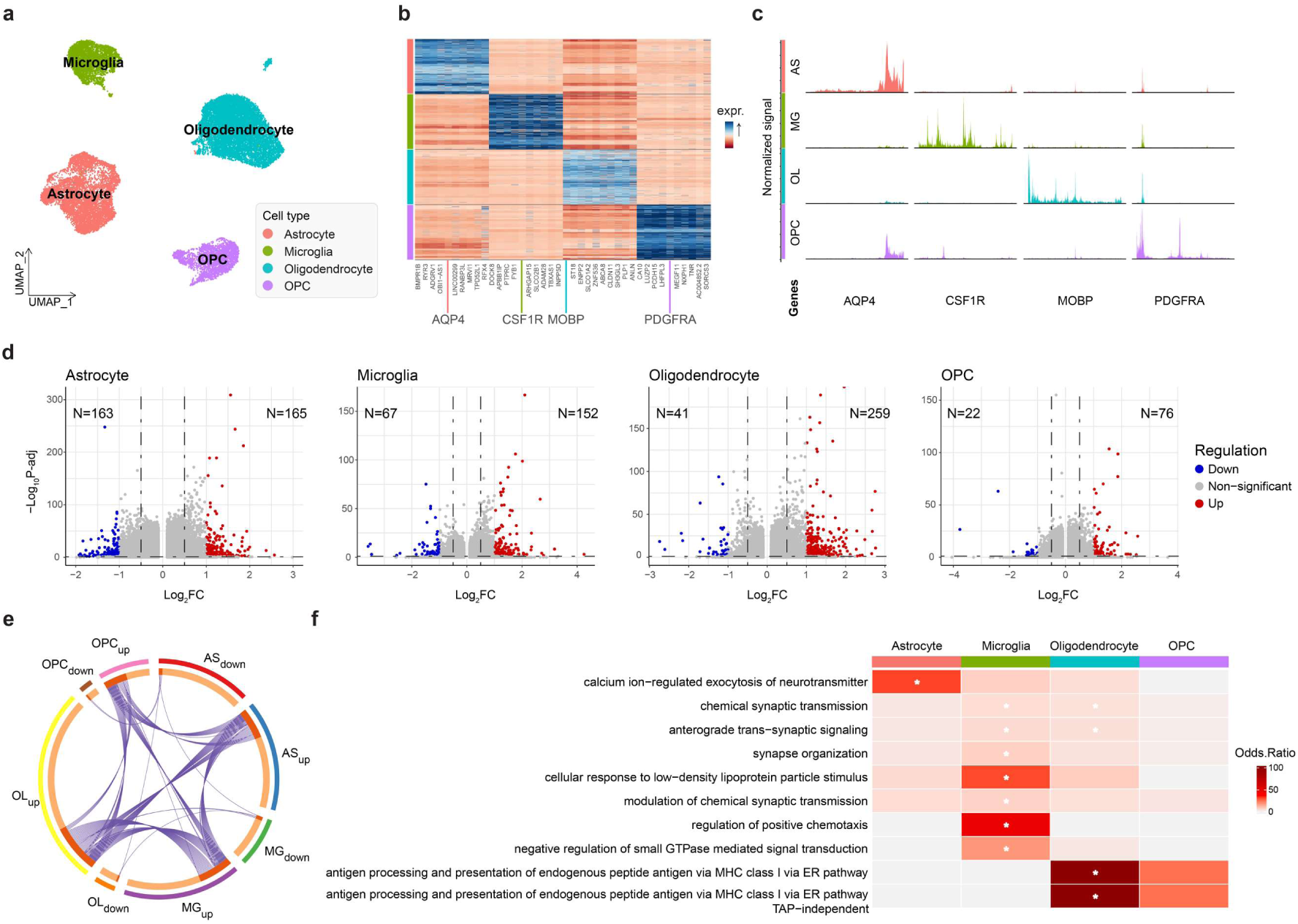
Cellular diversity and cell type-specific transcriptome dysregulation in Alzheimer’s middle temporal gyrus (MTG) revealed by glial subset of SEA-AD multiome data. (**a**) UMAP visualization of cell state space learned by scPair, with each color representing a distinct cell type (Astrocytes, Microglia, Oligodendrocytess, OPCs). (**b**) Row-normalized expression heatmap of cell type-specific markers based on the RNA component of SEA-AD multiome data, highlighting marker gene expression profiles across different cell types. (**c**) Chromatin accessibility across cell types visualized for major glial cell type marker genes highlighted in (b) based on the ATAC component of SEA-AD multiome data, illustrating accessible regions near marker genes in different cell populations. (**d**) Volcano plots showing differential gene expression (DEG) patterns in Alzheimer’s disease (AD) versus controls across major glial cell types. Points represent genes, with “down” (blue) indicating genes downregulated in AD, and “up” (red) for genes upregulated in AD. (**e**) Circos plot showing overlapping DEGs among different cell type groups, identifying shared dysregulated genes across cell populations. (**f**) Heatmap of the Odds ratio (which quantifies the association strength between each gene module and corresponding GO terms) for the top Gene Ontology (GO) terms enriched among upregulated DEGs within each cell type using enrichR. Blocks marked with asterisks represent adjusted *p* < 0.01, indicating significant enrichment.

We next performed differential gene expression (DEG) analysis in each glial cell type to investigate transcriptomic changes associated with AD. Compared to normal samples, we identified 165 upregulated and 163 downregulated genes in astrocyte, 152 upregulated and 67 downregulated genes in microglia, 259 upregulated and 41 downregulated genes in oligodendrocyte, and 76 upregulated and 22 downregulated genes in OPC (**Figure 3d**).

Furthermore, we compared these DEGs and found that many AD-upregulated genes were shared across multiple glial cell types, suggesting potentially common pathological mechanisms underlying glial dysfunction in AD (**Figure 3e**). To further explore the functional implications of these common genes, we performed pathway enrichment analysis across glial subtypes, and the results revealed distinct but interconnected pathways affected in each cell type. Astrocytes showed significant upregulation of calcium ion-regulated exocytosis of neurotransmitters pathway, which suggests potential alteration in neurotransmitter release and synaptic function. Microglia had the largest number of upregulated pathways, mostly related to synaptic functions, including chemical synaptic transmission, anterograde trans-synaptic signaling, synapse organization, cellular response to low-density lipoprotein particle stimulus, modulation of chemical synaptic transmission, regulation of positive chemotaxis, and negative regulation of small GTPase mediated signal transduction. Among these, chemical synaptic transmission and anterograde trans-synaptic signaling were also significantly upregulated in oligodendrocytes, in addition to oligodendrocytes’ specific antigen processing and presentation of endogenous peptide antigen via MHC class I via ER pathway, which indicates a possible role in immune signaling and neuroinflammation. OPCs also exhibited mild enrichment in those two antigen related pathways, though this did not reach statistical significance (**Figure 3f**).

### scMultiome analysis revealed key regulatory elements in in astrocytes

With glial sub-cell types identified, we first investigated key regulatory programs in astrocytes, which make up the majority of glial population in the human central nervous system. Astrocytes are essential regulators of glutamate and ion homeostasis, cholesterol and sphingolipid metabolism, and they respond dynamically to environmental factors, all of which have been implicated in neurological diseases, including AD (39). To further investigate astrocyte heterogeneity and its potential role in Alzheimer’s disease, we performed sub-clustering of astrocytes using scPair-generated embeddings to identify distinct astrocyte subpopulations.

This analysis revealed 13 astrocyte subclusters present in both AD and normal samples (**Figure 4a**). Interestingly, we observed significant differences in the proportion of these subclusters, with subclusters 5, 6, 7 and 11 predominantly enriched in AD patients (**Figure 4b**). To identify key regulators driving astrocyte dysfunction in AD, we computed Domains of Regulatory Chromatin (DORC) scores across these subclusters, and identified a total of 164 genes with high regulatory potential, hereafter referred to as DORC genes (**Figure 4c**). Next, we linked the transcription factors and their target genes by correlating the mRNA expression levels of transcription factors (TFs) with the accessibility of DORC-associated peaks. This TF-DORC regulatory network inference, built directly from the dataset rather than prior knowledge, served as another key step of our workflow and provided an unbiased representation of the gene-chromatin relationship in AD astrocytes, which identified both activating and repressive regulatory relationships (**Figure 4d**).

**Figure 4.**
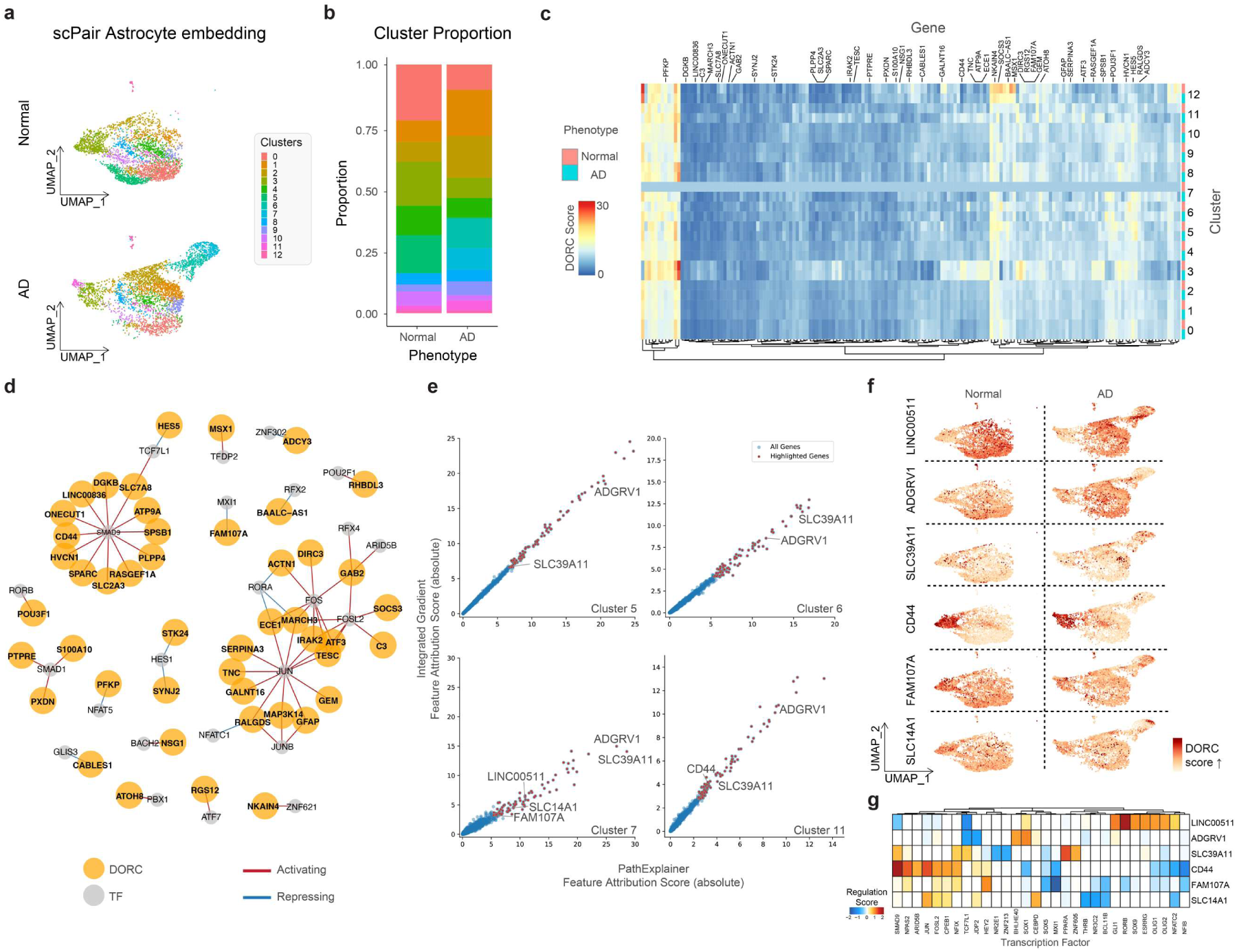
Integrative multi-Omics analysis reveals key regulatory modules in Alzheimer’s Disease Astrocytes. **(a)** UMAP visualization of astrocyte cell state space learned by scPair, with each color representing a distinct subpopulation. Top panel: normal control; Bottom panel: AD case. **(b)** Bar plot comparing the relative abundance of astrocyte subpopulations (clusters) between normal and AD groups. **(c)** Heatmap illustrating the Domains of Regulatory Chromatin (DORC) score all DORCs genes across conditions and cell clusters (n = 164). **(d)** TF-DORC network visualization for selected DORCs (orange nodes) and their associated TFs (gray nodes). Edges are scaled and colored by the signed regulation score. Red and blue edges indicate activating and repressive regulation, respectively. **(e)** Scatterplots showing the absolute feature attribution scores of genes calculated using the Path Explainer (x-axis) and Integrated Gradients (y-axis) methods on scPair model for corresponding clusters. The red-colored genes represent the intersection of the top 100 genes identified by both methods within each subpopulation (i.e. clusters 5, 6, 7, 11).. The displayed gene names are representatives of highly attributed genes intersecting with identified DORC genes. (**f**) UMAP visualization of astrocyte cell state space learned by scPair, split by selected intersecting genes and disease status. The color gradient represents the DORC score for each cell. Left: normal control. Right: AD case. (**g**) Heatmap of DORC regulation scores showing significant TF-DORC enrichments for selected DORC genes (*LINC00511, ADGRV1, SLC39A11, FAM107A, SLC14A1, CD44*) across clusters 5, 6, 7, 11. These genes are considered important based on absolute feature attribution scores in corresponding cluster(s) and have been linked to Alzheimer’s disease (AD) in published studies. The heatmap is colored by regulation scores.

We then applied two widely used feature attribution methods, Integrated Gradients (33) and Path Explainer (34), to rank genes based on their attribution scores through the scPair RNA-specific encoder network in a cell-subpopulation specific manner. Notably, this approach identified 15 genes that were both highly ranked by feature attribution (FA) analysis and identified as DORCs (**Figure 4e-f**), including *GFAP*, *LINC00511*, *TNC*, *SLC39A11*, *ADGRV1*, *CABLES1*, *FAM107A*, *DCLK1*, *LRIG1*, *HPSE2*, *CD44*, *SLC14A1*, *GALNT15*, *SPARCL1*, and *SERPINA3*. Given the overrepresentation of subclusters 5, 6, 7 and 11 in AD, we specifically investigated the DORC and FA genes enriched within these clusters, and identified *ADGRV1*, *SLC39A11*, *SLC14A1*, *FAM107A*, *CD44* and *LINC00511* as potential key elements in AD-related astrocyte regulation and dysfunction (**Figure 4e**).

*ADGRV1* (Adhesion G Protein-Coupled Receptor V1) is a crucial regulator of glutamate uptake and regulates homeostasis in astrocytes, and it plays key roles in nervous system development including cortical patterning, dendrite and synapse formation, and myelination (40). *SLC39A11* and *SLC14A1* encode Solute Carrier proteins. *SLC39A11* has been patented as a potential diagnostic and therapeutic target for AD (41). *SLC14A1*, also known as *Urea Transporter B* (*UT-B*), is the primary transporter of urea in the brain and has been reported to be downregulated in AD astrocytes. Therapeutic targets that increases *SLC14A1* expression, such as *Gene-silencing of astrocytic ornithine decarboxylase-1* (*ODC1*), has been shown to help the recovery from reactive astrogliosis and memory impairment in AD model (42). *FAM107A* encodes Family With Sequence Similarity 107 Member A. It is involved in cognitive functions and has been classified as a tumor suppressor gene (43,44), while recent studies suggest it may have immune-related functions relevant to AD (45,46). *CD44*, a major surface hyaluronan (HA) receptor, mediates cell adhesion, migration and signaling (47). In Alzheimer’s disease brain, the number of *CD44*-positive astrocytes increased dramatically, indicating a potential role in neuroinflammation and gliosis (48). *LINC00511* is a long non-coding RNA that has been identified as a prognostic biomarker in multiple cancers including breast cancer, ovarian cancer, liver cancer, pancreatic cancer, lung cancer and glioma (49). While no direct associations to AD have been reported yet, its established negative regulatory role in cell proliferation, cell cycle progression, apoptosis, invasion, and migration (50–52) suggest a potential involvement in AD-associated astrocyte dysfunction and AD pathogenesis.

Finally, our comprehensive analysis also highlighted important AD-related transcription factors in astrocytes. For instance, *JUN* and *FOSL2* are well-known transcription factors in Alzheimer’s disease (AD) research, particularly in the context of astrocytes. They are part of the transcription factor activator protein 1 (AP-1) (53) family, which is involved in a variety of important cellular processes in AD such as inflammation, stress response, and cell differentiation (**Figure 4g**). *FOSL2*, specifically, has been reported to activate the disease-associated astrocyte signature (54). *PPARA* is another important AD-related transcription factor. It regulates the transcription of genes related to the metabolism of cholesterol, fatty acids, other lipids and neurotransmission, mitochondria biogenesis (55). *PPARA* has a profound relationship with *APP*, one of the major genetic risk factors of AD. In fact, it has been reported that administration of *PPARA* agonists decreases amyloid pathology and reverses memory deficits and anxiety symptoms in *APP*-*PSEN1* mouse model (56). Our study’s capturing of these key regulator of AD demonstrated the utility of our workflow to nominate therapeutic targets through interpretable machine learning.

### scMultiome analysis uncovers an AD-specific state in microglia

We next focused on microglia, the brain’s primary immune cells, which play a crucial role in AD by transitioning from homeostatic to activated states throughout disease progression (57) (58). To gain deeper insights into the mechanisms underlying this heterogeneity, we performed sub-clustering using scPair-derived cell embeddings and identified 11 distinct microglial subpopulations shared between AD cases and controls (**Figure 5a**). The proportions of these subclusters were calculated (**Figure 5b**), revealing differences between the two groups and suggesting AD-associated shifts in microglial phenotypes.

**Figure 5.**
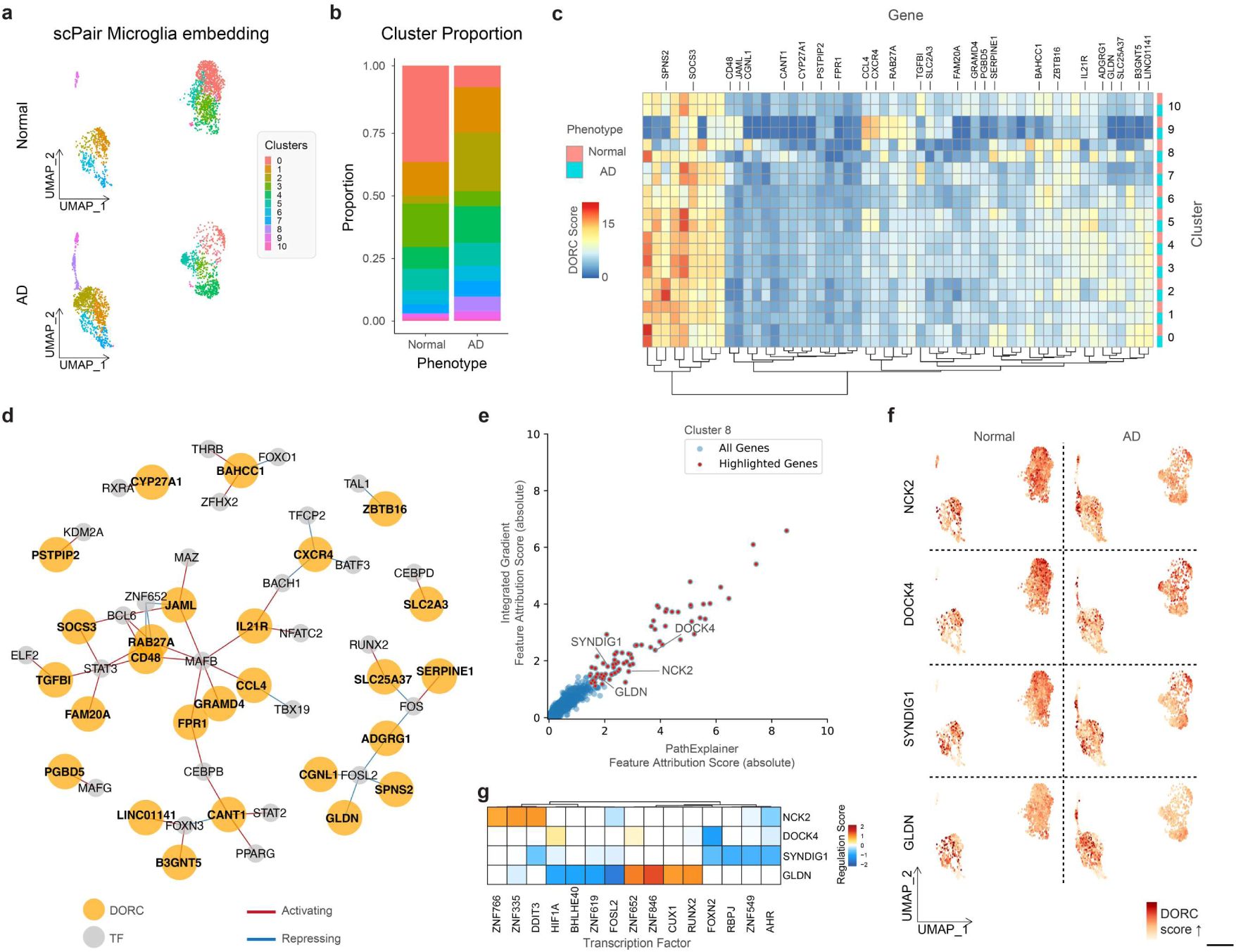
Integrative multi-Omics analysis reveals key regulatory elements in Alzheimer’s Disease Microglia. **(a)** UMAP visualization of microglial cell state space learned by scPair, with each color representing a distinct subpopulation. Top panel: normal control; Bottom panel: AD case. **(b)** Bar plot comparing the relative abundance of microglial subpopulations (clusters) between normal and AD groups. **(c)** Heatmap illustrating the Domains of Regulatory Chromatin (DORC) score all DORCs genes across conditions and cell clusters (n = 56). **(d)** TF-DORC network visualization for selected DORCs (orange nodes) and their associated TFs (gray nodes). Edges are scaled and colored by the signed regulation score. Red and blue edges indicate activating and repressive regulation, respectively. **(e)** Scatterplots showing the absolute feature attribution scores of genes calculated using the Path Explainer (x-axis) and Integrated Gradients (y-axis) methods on scPair model for corresponding cluster(s). The red-colored genes represent the intersection of the top 100 genes identified by both methods within each subpopulation (i.e. cluster 8). The displayed gene names are representatives of highly attributed genes intersecting with identified DORC genes. (**f**) UMAP visualization of microglial cell state space learned by scPair, split by selected intersecting genes and disease status. The color gradient represents the DORC score for each cell. Left: normal control. Right: AD case. **(g)** Heatmap of DORC regulation scores showing significant TF-DORC enrichments for selected DORC genes (*NCK2, DOCK4, SYNDIG1*, and *GLDN*) in cluster 8. These genes are considered important based on absolute feature attribution scores in corresponding cluster(s) and have been linked to Alzheimer’s disease (AD) in published studies. The heatmap is colored by regulation scores.

Further exploration of the regulatory landscape of microglia subtypes was conducted using DORC and FA analysis. As a result, a TF-DORC regulatory network for microglia was constructed with 56 DORC genes (**Figure 5c-d**). 14 of these genes were highly ranked in FA analysis, including *SLC11A2, SMAP2, SYNDIG1, DENND3, NCK2, MRC1, ZBTB16, GLDN,*

*ACSL1, DOCK4, PALD1, RGS1, TMSB4X*, and *SRGN*. We found that among all the microglia subtypes we identified, there was a subcluster nearly unique to AD cases. This AD-specific microglia subcluster, cluster 8 (**Fig. 5a, b**), was almost exclusively composed of AD microglia, indicating that it may be uniquely driven by AD pathology. Within this cluster, four DORC genes, *NCK2*, *DOCK4*, *SYNDIG1*, and *GLDN* exhibited high feature attribution scores, suggesting that they can be key regulators in AD pathology under this particular cell state (**Figure 5e-f**).

*NCK2* encodes NCK Adaptor Protein 2 and was recently reported to be highly expressed in amyloid-responsive microglial, and a newly-discovered rare variant in *NCK2* has also been linked to late-onset AD (59). *DOCK4,* encoding Dedicator Of Cytokinesis 4, has been associated with neuropsychiatric diseases such as autism (60), dyslexia (61), and schizophrenia (62), but a recent multi-trait association study linked a low-frequency coding variant of *DOCK4* to known AD loci (63). *SYNDIG1*, which encodes Synapse Differentiation Inducing 1, belongs to the interferon-induced transmembrane family and regulates excitatory synapse number and strength in hippocampal neurons (64). A genome wide association study (GWAS) further identified *SYNDIG1* as one of the loci significantly associated with Braak staging, a measure of AD severity (65). *GLDN*, encoding Gliomedin, functions as a glial ligand for neurofascin and NrCAM, two axonal immunoglobulin cell adhesion molecules critical for organizing sodium channels at the nodes of Ranvier. While its role in peripheral nervous system node formation is established, its involvement in AD remains undocumented (66), making it a potential promising target for further investigation.

In addition to these important regulatory genes, our detailed analysis also revealed key transcription factors associated with Alzheimer’s disease in microglia. (**Figure 5g**). Interestingly, *ZNF766, ZNF335, ZNF619, ZNF652, ZNF846*, and *ZNF549* from the Zinc Finger (ZNF) family were enriched in cluster 8, the AD-specific microglia subcluster. Several ZNF family genes have been implicated in amyloid-β (Aβ) deposition and neurofibrillary tangle accumulation, the two major hallmarks of AD (67). Another TF of interest was *HIF-1α*, a key regulator of cellular and tissue adaption to low oxygen tension. *HIF-1α* has gained attention as a potential medicinal target for neurodegenerative disease due to its dual role in microglia regulation: it can inactivate microglia and cause microglia death and promotes neuroinflammation; on the other hand, it can inhibit tau hyperphosphorylation and promote microglial activation (68) (69).

### scMultiome profiling uncovers new insights into regulatory drivers in oligodendrocytes and OPCs

The last major glial cell populations are oligodendrocyte and oligodendrocyte progenitor cells (OPCs), and we analyzed their transcriptional and chromatin accessibility profiles in both control and AD. Oligodendrocytes are responsible for producing myelin, a specialized elongated membrane structure that tightly wraps around axons to provide insulation and metabolic support (70). OPCs serve as their progenitors, which plays crucial role in both myelin formation and repair (71). Dysregulation of oligodendrocytes and OPCs, and the subsequent demyelination process are recognized as key pathobiological events in AD.

After re-clustering the previously defined oligodendrocytes and OPCs using scPair embedding, we observed 11 subpopulations present in both AD cases and controls (**Figure 6a-b**). A total of 142 DORC genes with high regulatory potential were identified (**Figure 6c**), and a TF-DORC network was constructed to map key regulatory interactions (**Figure 6d**). To further refine these findings, feature attribution overlapping analysis was applied, leading to the identification of five top-scoring DORC genes (**Figure 6e-f**): *RASGRF2, BEST3, SLC8A1, SLC5A11*, and *SNTG1*. Among these, *SLC8A1* and *SNTG1* were predominantly expressed in clusters 3 and 9, which showed significant enrichment in AD patients (**Figure 6b)**. *SLC8A1*, which encodes sodium/calcium exchanger 1 (NCX1), plays a critical role in maintaining intracellular Na^+^ and Ca2^+^ homeostasis under pathophysiological conditions in the brain [30].

**Figure 6.**
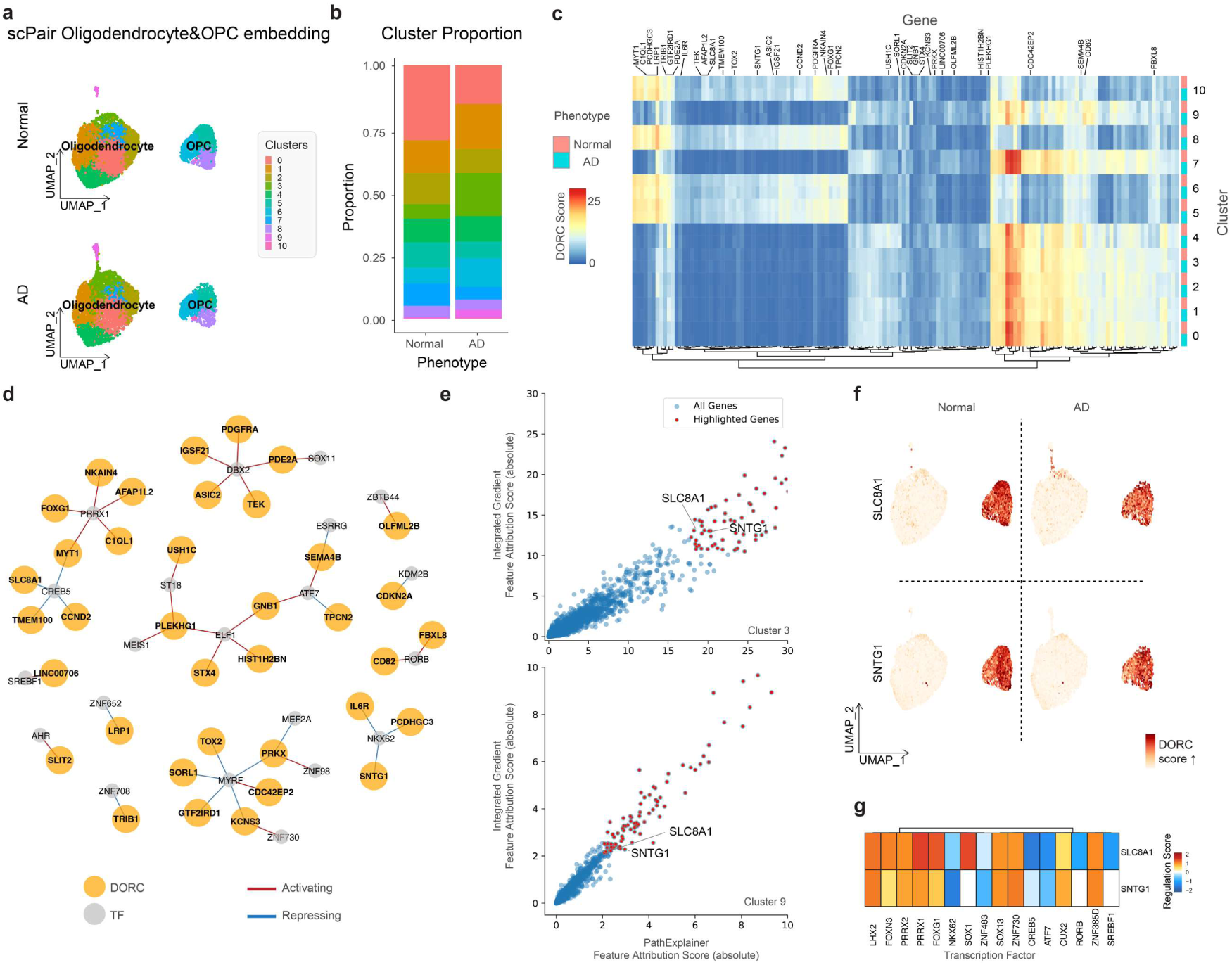
Integrative multi-Omics analysis reveals key regulatory modules in Alzheimer’s Disease Oligodendrocytes and OPC. **(a)** UMAP visualization of oligodendrocytes and OPC cell state space learned by scPair, with each color representing a distinct subpopulation. Top panel: normal control; Bottom panel: AD case. **(b)** Bar plot comparing the relative abundance of microglial subpopulations (clusters) between normal and AD groups. **(c)** Heatmap illustrating the Domains of Regulatory Chromatin (DORC) score all DORCs genes across conditions and cell clusters (n = 142). **(d)** TF-DORC network visualization for selected DORCs (orange nodes) and their associated TFs (gray nodes). Edges are scaled and colored by the signed regulation score. Red and blue edges indicate activating and repressive regulation, respectively. **(e)** Scatterplots showing the absolute feature attribution scores of genes calculated using the Path Explainer (x-axis) and Integrated Gradients (y-axis) methods on scPair model for corresponding cluster(s). The red-colored genes represent the intersection of the top 100 genes identified by both methods within each subpopulation (i.e. clusters 3 and 9). The displayed gene names are representatives of highly attributed genes intersecting with identified DORC genes. (**g**) UMAP visualization of oligodendrocytes and OPC cell state space learned by scPair, split by selected intersecting genes and disease status. The color gradient represents the DORC score for each cell. Left: normal control. Right: AD case. **(f)** Heatmap of DORC regulation scores showing significant TF-DORC enrichments for selected DORC genes (*SLC8A1* and *SNTG1*) across clusters 3 and 9. These genes are considered important based on absolute feature attribution scores in corresponding cluster(s) and have been linked to Alzheimer’s disease (AD) in published studies. The heatmap is colored by regulation scores.

While a related family member NCX3 has been showed to be essential for oligodendrocyte differentiation [31], our study is the first to link *SLC8A1* to oligodendrocyte dysregulation in AD. *SNTG1* encodes Syntrophin Gamma 1 and plays an important role in synaptic organization, transmission, and cognitive function. It has previously been identified as a potential biomarker in dementia [34], and our findings suggest that its involvement in AD progressions may be regulated in oligodendrocytes.

Apart from these candidate DORC genes, our analysis identified TFs with potential regulatory influence in oligodendrocyte-lineage cells in AD (**Figure 6g**). *PRRX1* and *PRRX2*, transcriptional co-activators, were previously highlighted as active regulons in OPCs in AD through gene regulatory network analysis (72). We also identified *SOX1* and *SOX13*, members of the SOX family of transcription factors, which not only function as classical TFs but also serve architectural roles in chromatin remodeling. These factors are traditionally associated with oligodendrocyte myelination (73). Emerging research has begun to reveal their involvement in pathological conditions in past years. For instance, a recent transgenic mouse model study reported significant differences in *SOX1*-positive cell populations between disease and control groups (74). Finally, we identified *SREBF1*, a master regulator of cholesterol homeostasis that is often inactivated by Aβ. Interestingly, a recent single-cell multiome study reported downregulation of SREBF1 and its target genes specifically in AD oligodendrocytes (54), which further supports its relevance to AD pathogenesis.

## DISCUSSION

scMultiome offers a powerful approach to investigate cellular diversity and disease mechanisms. In this study, we built a robust analytical workflow for integrative analysis of paired scRNA-seq and scATAC-seq data, addressing key challenges such as effective batch correction, cross-modality alignment and prediction. By applying out workflow to the SEA-AD Consortium’s single-cell multiome dataset, we identified regulatory programs mediated by key transcription factors in astrocytes, microglia, and cells in oligodendrocytes lineages, shedding light on their roles in Alzheimer’s disease pathogenesis. Our findings revealed AD-associated shifts in glial subpopulations, identified transcriptional and epigenomic signatures, and established regulatory networks linking critical genes to transcription factors. Furthermore, we applied interpretable machine learning techniques to refine our understanding of regulatory programs, pinpointing genes with high contributing values in disease-related chromatin accessibility prediction.

During our benchmarking, we highlighted the advantages of deep learning-based approaches over traditional factorization-based methods, and demonstrated that capturing complex multimodal interplay or relationships worked better in a supervised manner compared to unsupervised methods which heavily relied on highly variable features. Traditionally, interpretability of deep learning approaches is challenge because embeddings learned by deep learning models do not inherently reflect biological meaning (34). To bridge this gap, we employed feature attribution techniques, allowing us to extract biologically relevant insights from the learned representations, which worked especially well on end-to-end models such as scPair. By comparing feature attribution-derived genes with DORC genes, an alternative multimodal correlation-based functional inference method, we identified overlapped key drivers of glial regulatory changes in AD, demonstrating the utility of this approach for inferring mechanistic insights from high-dimensional single-cell multiomic data.

The majority of AD cases involve a complex interplay of genetics and environmental factors, with the contribution of many genes yet to be discovered. Our study utilized an integrated explainable machine learning and multiomics approach to identify novel and support known key genes that have high regulatory potential in AD pathogenesis. Among these, we discovered several genes in the SLC family, including *SLC39A11* and *SLC14A1* in astrocytes and *SLC8A1* and *SLC5A11* in oligodendrocytes. The SLC family is a large group of transport proteins that play a crucial role in moving various substances, including ions, neurotransmitters, and nutrients, across cell membranes. While some genes in the SLC family are known to be associated with neurological disorders, such as *SLC2A1*, *SLC22A8, and SLCO1A2* (*75–77*), there has been no direct link between SLC family genes to AD. Our study is the first to name SLC family genes as key regulatory genes in AD in multiple cell types, highlighting the application of our approach to identifying target genes and regulatory networks that may otherwise be overlooked.

The importance of glial cells in AD pathogenesis has been gaining increasing recognition in recent years. Indeed, unveiling the intricate functions of glial cells may open new avenues for understanding the underlying mechanisms of neurodegeneration and identifying potential therapeutic targets. When re-clustering glial cells, we found cell subtypes that were particularly affected by AD, such as subclusters 5, 6, 7, and 11 in astrocyte, subcluster 8 in microglia, and subclusters 3 and 9 in oligodendrocyte-lineage. Prior studies have reported that astrocytes, microglia, and oligodendrocytes undergo subtype-specific transcriptional changes in Alzheimer’s disease (78–80), and our findings not only recapitulated these shifts but also provided new insights into the regulatory programs underlying these changes. The identification of both known and novel driver genes in AD-associated glial subpopulations suggests that distinct molecular mechanisms contribute to glial dysfunction in different cellular contexts. The integration of regulatory network inference with interpretable machine learning further strengthens the translational relevance of our findings.

Moving forward, the application of our workflow to additional brain regions and disease contexts may further expand our understanding of glial regulatory dynamics in neurodegeneration. Moreover, integrating other single-cell modalities, such as spatial transcriptomics and proteomics, could provide additional layers of resolution for dissecting cell-cell interactions and disease-associated microenvironments. As single-cell multiomic technologies continue to evolve, incorporating advanced machine learning techniques alongside biological domain knowledge will be essential for unlocking the full potential of these datasets.

## DATA AVAILABILITY

The multiome data used in this study can be accessed from AD knowledge Portal under ID: syn29879329: https://www.synapse.org/Synapse:syn29879329

## AUTHOR CONTRIBUTIONS

S.Z. obtained, preprocessed the multiome data, and designed the workflow. H.H. and S.Z. conducted the benchmarking of integration methods. S.Z. performed formal analysis and the case studies. Y.W.A. and Y.R. supervised the project. S.Z., H.H. and Y.R. wrote the manuscript with reviews from X.W., C.X., and Y.W.A.

## ACKNOWLEDGEMENTS

Data used in this study were obtained from the Seattle Alzheimer’s Disease Brain Cell Atlas (SEA-AD) consortium. The work presented in the study is supported by U19 AG069701 to C.X.,

Y.W.A. and Y.R., and Mayo Clinic Department of Quantitative Health Sciences Associate Consult fund to Y.R. and X.W.

## FUNDING

This work is supported by U19 AG069701 to C.X., Y.W.A. and Y.R., and Mayo Clinic Department of Quantitative Health Sciences Associate Consult fund to Y.R. and X.W..

## CONFLICT OF INTEREST

The authors declare no conflict of interest.

## Notes

### Competing Interest Statement

The authors have declared no competing interest.

